## Supplementary material for "Metagenomics uncovers dietary adaptations for chitin digestion in the gut microbiota of convergent myrmecophagous mammals": Figure S1

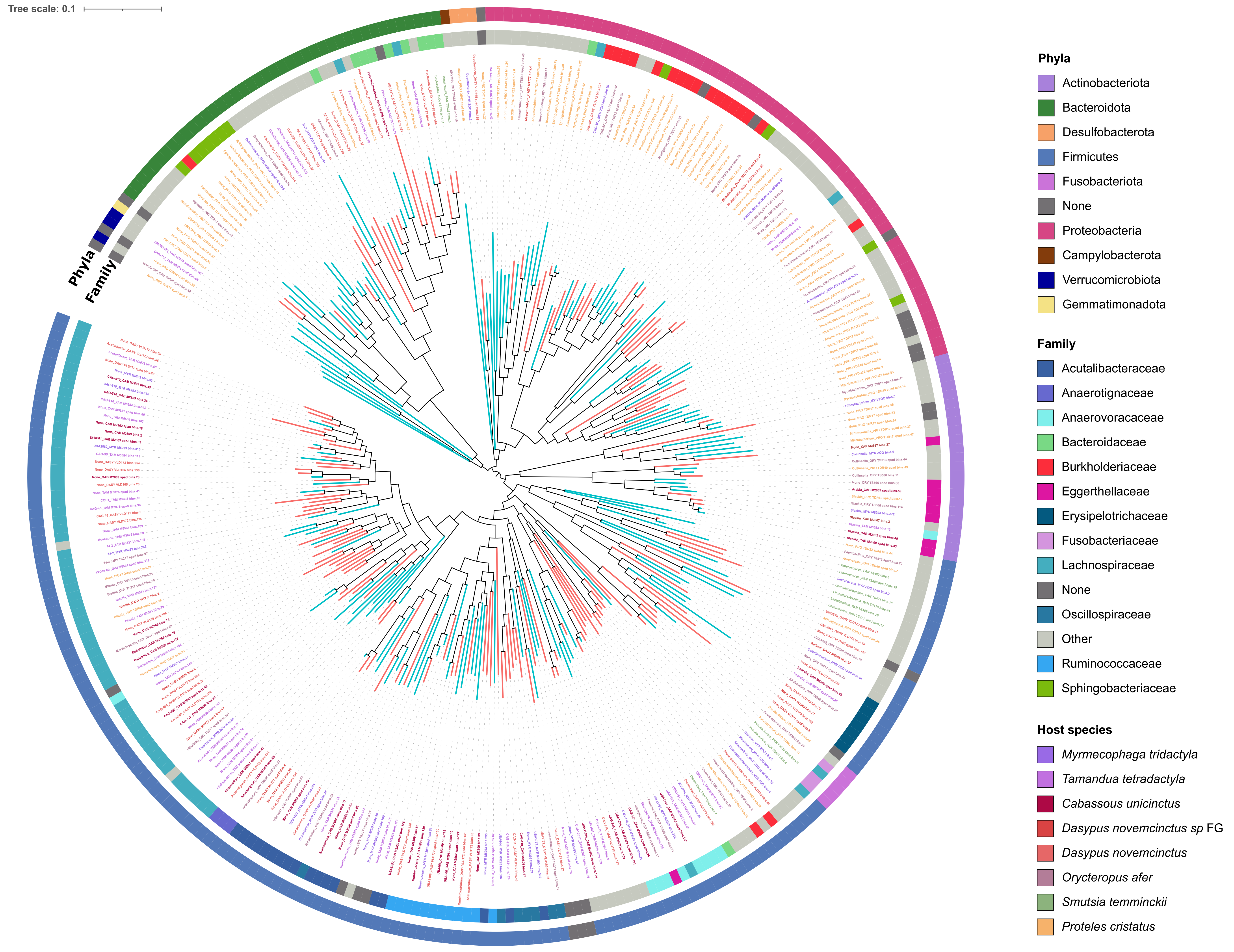


**Fig S1 Phylogeny of the 314 high-quality selected bins reconstructed from long read assemblies** (n = 156; red branches) **and short-read assemblies** (n = 158; blue branches)**.** Circles respectively indicate (from inner to outer circles): the bacterial family and phyla the bin was assigned to based on the Genome Taxonomy Database release 7 (38). Colored sequence names indicate the host species. Bins' names of the myrmecophagous-specific clades are indicated at leaves of the phylogenetic tree together with the genus to which they were assigned to.
