## Supplementary material for "Metagenomics uncovers dietary adaptations for chitin digestion in the gut microbiota of convergent myrmecophagous mammals": Figure S2 A

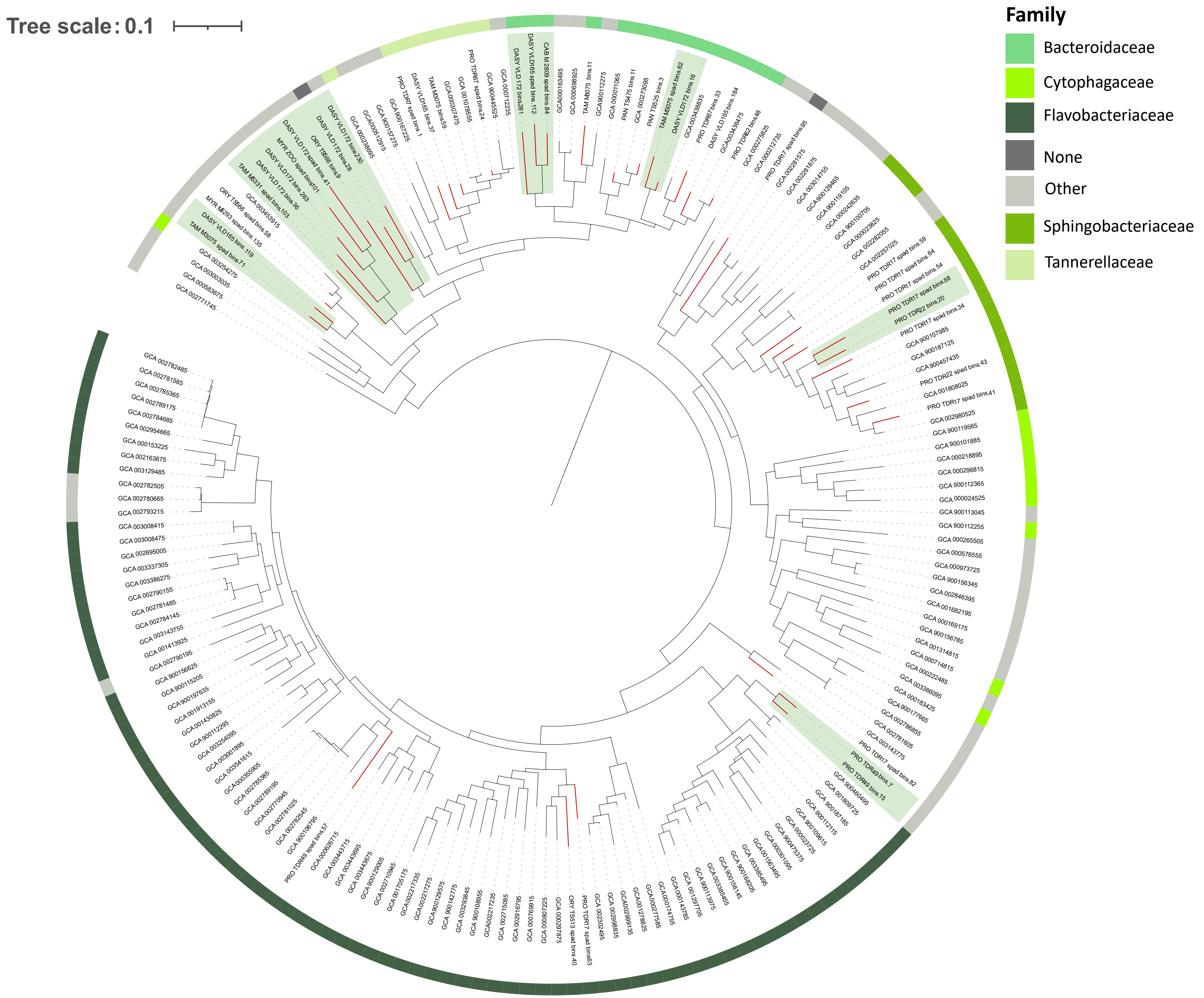


**Fig S2 A Myrmecophagous-specific clades within Bacteroidetes** (green highlights).

This tree is a subtree of Fig. 1. Outer circles indicate the bacterial family to which these genome bins were assigned based on the Genome Taxonomy Database release 7 (38).
