## Supplementary material for "Metagenomics uncovers dietary adaptations for chitin digestion in the gut microbiota of convergent myrmecophagous mammals": Figure S2 B

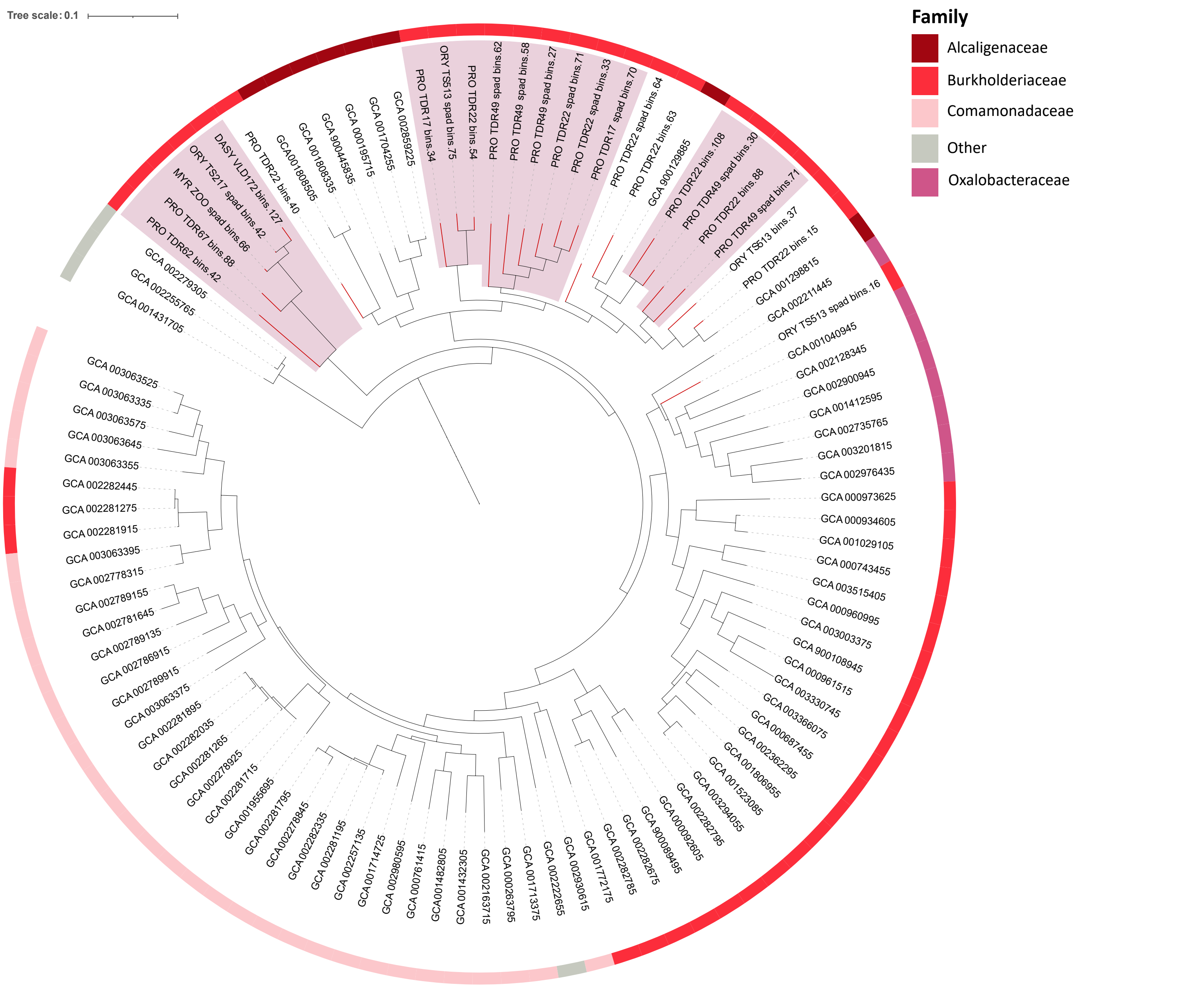


**Fig S2 B Myrmecophagous-specific clades within Proteobacteria** (red highlights).
This tree is a subtree of Fig. 1. Outer circles indicate the bacterial family to which these genome bins were assigned based on the Genome Taxonomy Database release 7 (38).
