## Supplementary material for "Metagenomics uncovers dietary adaptations for chitin digestion in the gut microbiota of convergent myrmecophagous mammals": Figure S3

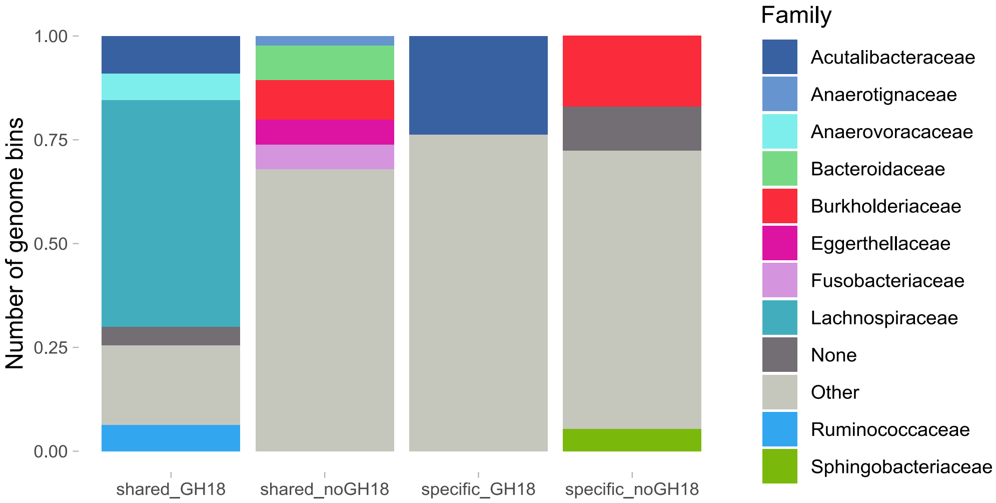


**Fig S3 Bacterial families of host-species specific and shared genome bins carrying or not carrying GH18 genes.**
