## Supplementary material for "Metagenomics uncovers dietary adaptations for chitin digestion in the gut microbiota of convergent myrmecophagous mammals": Figure S4

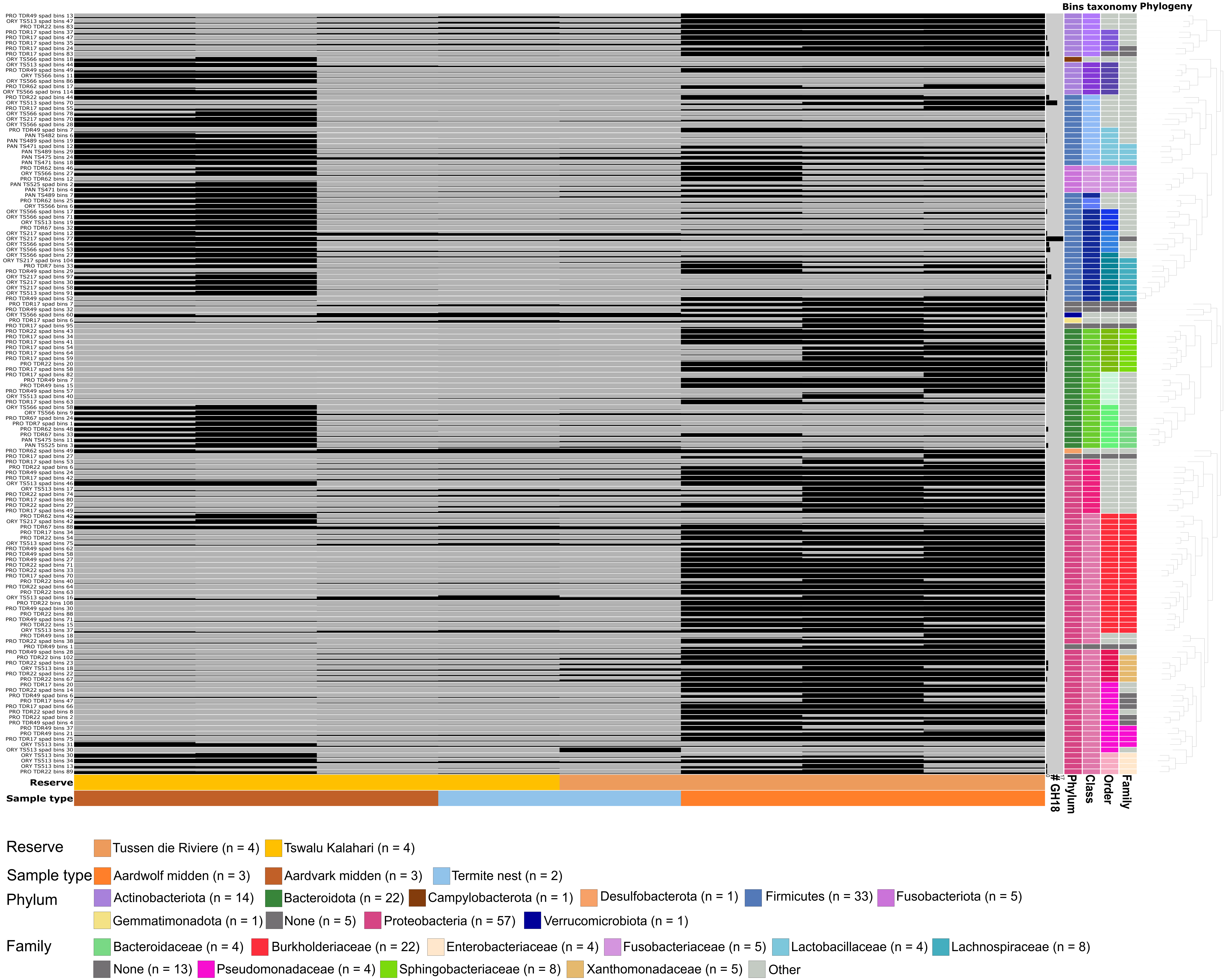


**Fig S4 Detection of the 140 high-quality selected bins reconstructed from the aardvark, ground pangolin, and southern aardwolf samples** (lines) **in the eight soil samples** (columns), **collected in South Africa near feces sampling sites.** Each square indicates the detection of a bin in a sample as estimated by anvi'o v7 (41). Bin names are indicated on the left. The provenance of each sample is indicated by different colors at the bottom of the graph. Columns on the right indicate (from left to right): the number of GH18 sequences identified in each bin (from 0 to 17), the bin's t axonomic phylum, class, order, and family. The phylogeny of the 140 selected bins inferred with PhyloPhlAn v3.0.58 (42) is also represented on the right of the graph. See detection table available via Zenodo for detailed values of detection.
