## Supplementary material for "Metagenomics uncovers dietary adaptations for chitin digestion in the gut microbiota of convergent myrmecophagous mammals": Table S3

**Table S3 Proportion of chitinolytic genome bins** (genomes having at least one GH18 with an active chitinolytic site) **detected in the nine focal myrmecophagous species.**

| Species | # chitinolytic bins detected | total # detected bins | # chitinolytic bins / total # bins detected |
|---|---|---|---|
| ***Cabassous unicinctus*** | 46 | 113 | 0.407079646 |
| ***Dasypus sp. nov.* FG** | 48 | 105 | 0.457142857 |
| ***Dasypus novemcinctus*** | 50 | 129 | 0.387596899 |
| ***Dasypus kappleri*** | 2 | 25 | 0.08 |
| ***Myrmecophaga tridactyla*** | 48 | 141 | 0.340425532 |
| ***Orycteropus afer*** | 15 | 97 | 0.154639175 |
| ***Smutsia temminckii*** | 5 | 18 | 0.277777778 |
| ***Proteles cristatus*** | 12 | 104 | 0.115384615 |
| ***Tamandua tetradactyla*** | 37 | 121 | 0.305785124 |
