## Supplementary material for "Metagenomics uncovers dietary adaptations for chitin digestion in the gut microbiota of convergent myrmecophagous mammals": Table S4

**Table S4 Detailed sample information for the eight soil samples collected in South Africa.**

| Sample | Sample type | Species | Common name | Class | Order | Family | Location |
|---|---|---|---|---|---|---|---|
| **TDR012** | Soil from Midden #1 | *Proteles cristatus* | Southern ardwolf | Mammalia | Carnivora | Hyaenidae | Tussen Die Riviere Reserve |
| **TDR014** | Nest fragments and soil | *Trinervitermes sp.* | Snouted termites | Insecta | Blattodea | Termitidae | Tussen Die Riviere Reserve |
| **TDR019** | Soil from Midden #2 | *Proteles cristatus* | Southern ardwolf | Mammalia | Carnivora | Hyaenidae | Tussen Die Riviere Reserve |
| **TDR023** | Soil from Midden #3 | *Proteles cristatus* | Southern ardwolf | Mammalia | Carnivora | Hyaenidae | Tussen Die Riviere Reserve |
| **TS218** | Soil from midden | *Orycteropus afer* | Aardvark | Mammalia | Tubulidentata | Orycteropodidae | Tswalu Kalahari Reserve |
| **TS274** | Nest fragments and soil | *Trinervitermes sp.* | Snouted termites | Insecta | Blattodea | Termitidae | Tswalu Kalahari Reserve |
| **TS281** | Soil from midden | *Orycteropus afer* | Aardvark | Mammalia | Tubulidentata | Orycteropodidae | Tswalu Kalahari Reserve |
| **TS298** | Soil from midden | *Orycteropus afer* | Aardvark | Mammalia | Tubulidentata | Orycteropodidae | Tswalu Kalahari Reserve |
